# Listening to music modulates visual perception of paintings during binocular rivalry

**DOI:** 10.1101/2024.09.17.613593

**Authors:** Xintong Jiang, Yuhui Cheng, Yi Jiang, Yi Du

## Abstract

Music, as a hierarchically organized art form, profoundly influences perception and mood through its rich information and intense emotion. While previous research has demonstrated that music can shape visual perception, its influences on visual awareness in aesthetic scenes remain largely unclear. In a series of five experiments involving 138 participants, we used the binocular rivalry paradigm to investigate how sad or lively Chinese folk music influences the perception of traditional Chinese paintings with varying aesthetic emotion (bleak vs. vivid). Paintings were presented in upright or inverted orientations or replaced by patches. Results revealed that sad music significantly increased both the average dominance duration and the predominance score of bleak paintings. Conversely, lively music with a faster tempo accelerated the switch rate of both paintings and patches. These effects were interdependent, exclusively for meaningful (upright) paintings, and correlated with individual sensitivity to musical reward. Our findings support the hierarchical predictive coding model of binocular rivalry, demonstrating that musical aesthetic emotion exerts top-down modulation, while musical tempo provides bottom-up entrainment, jointly shaping visual awareness of paintings.

## Introduction

Music, a ubiquitous art form, conveys rich emotions and meanings (1), influencing our perception and behavior (2,3). It affects our eye movements (4,5) and aesthetic experience (6) in natural scenes, and even disambiguates visual perception (7). But can music also shape our appreciation and perception of visual art? Imagine visiting a gallery, confronted with a dazzling, abstract painting: could background music guide your interpretation of the scene? The question remains unanswered, largely due to limited research on music’s role in multisensory aesthetic experiences, which may stem from the abstract and hierarchical nature of musical meaning.

The meaning of music operates on multiple levels: iconic meaning, derived from low-level properties like pitch, rhythm, and timbre, and indexical meaning, associated with musical emotion. Unlike discrete basic emotions (e.g., happiness, anger, fear), music emotions are considered as more “aesthetic” rather than “utilitarian” (8). Aesthetic emotions are complex and often inexpressible, triggered by the instinct of knowledge acquisition in higher animals, thus making them central to art (9). These emotions span a broader semantic range (10), and are represented at higher levels of mental representation (9). However, the interaction of aesthetic emotions between visual and auditory art forms remains largely underexplored.

Recent studies have attempted to examine the effect of music on visual perception using binocular rivalry, where conflicting monocular images are presented to each eye, causing spontaneously alternates in perceptual awareness between the two images (11). This paradigm allows researchers to separate visual input from perceptual awareness (12). Evidence has accumulated that visual features, such as stimulus frequency (13,14), orientation (13,15,16), and semantic content (17–19) can be cross-modally biased. However, previous research has primarily focused on the effects of concrete musical elements, such as basic emotions (20,21) and symbolic representation (22,23), on non-aesthetic aspects of visual perception. For instance, emotional music prolonges the perceived duration of congruent (happy) face compared to incongruent or neutral faces (20). Additionally, only individuals capable of read music (22) or those with absolute pitch (23) could recognize the congruence between melodic sound and musical scores.

So far, the influence of abstract, hierarchical representations of music on visual art perception in binocular rivalry remains unexplored, leaving the underlying mechanism unclear. We hypothesized that music modulates rivalry dynamics through both top-down and bottom-up processes. Aesthetic emotion of music may evoke specific feelings, serving as a contextual factor that biases the perceived content of paintings (top-down modulation), while musical tempo may drive visual fluctuations through bottom-up entrainment. Moreover, these top-down and bottom-up effects are likely to be interdependent. Additionally, aesthetic emotion elicited by artworks recruit the neural circuitry involved in reward (9). Although musical sensitivity has not been shown to differentiate aesthetic judgments of visual art and music (24), individuals with higher musical reward sensitivity are likely to exhibit a greater musical impact on binocular rivalry.

To test these hypotheses, we conducted five experiments. In Experiments 1 and 2, we created an artistic environment combining music and paintings to investigate the top-down effect of music’s aesthetic emotion on visual perception. In Experiment 3, we presented inverted paintings, whose aesthetic emotions are harder to recognize, alongside the upright paintings to further validate the top-down effect. Participants listened to traditional Chinese music (lively or sad) played by different instruments (erhu or bamboo flute) or no music at all, while viewing traditional Chinese paintings (vivid vs bleak) in a binocular rivalry task (see **Figure 1a** and **Procedure** in **Materials and Methods** for details). Participants confirmed the aesthetic emotions of music and paintings both in the pretest and after the binocular rivalry task (see **Figure 1c** and **Subjective assessments** in **Materials and Methods** for details). To elucidate the bottom-up effect, we transformed the paintings into patches and manipulated the musical tempo in Experiments 4 and 5. We also explored the correlations between the observed musical effects in binocular rivalry and their relationships with individuals’ sensitivity to musical reward.

**Figure 1.**
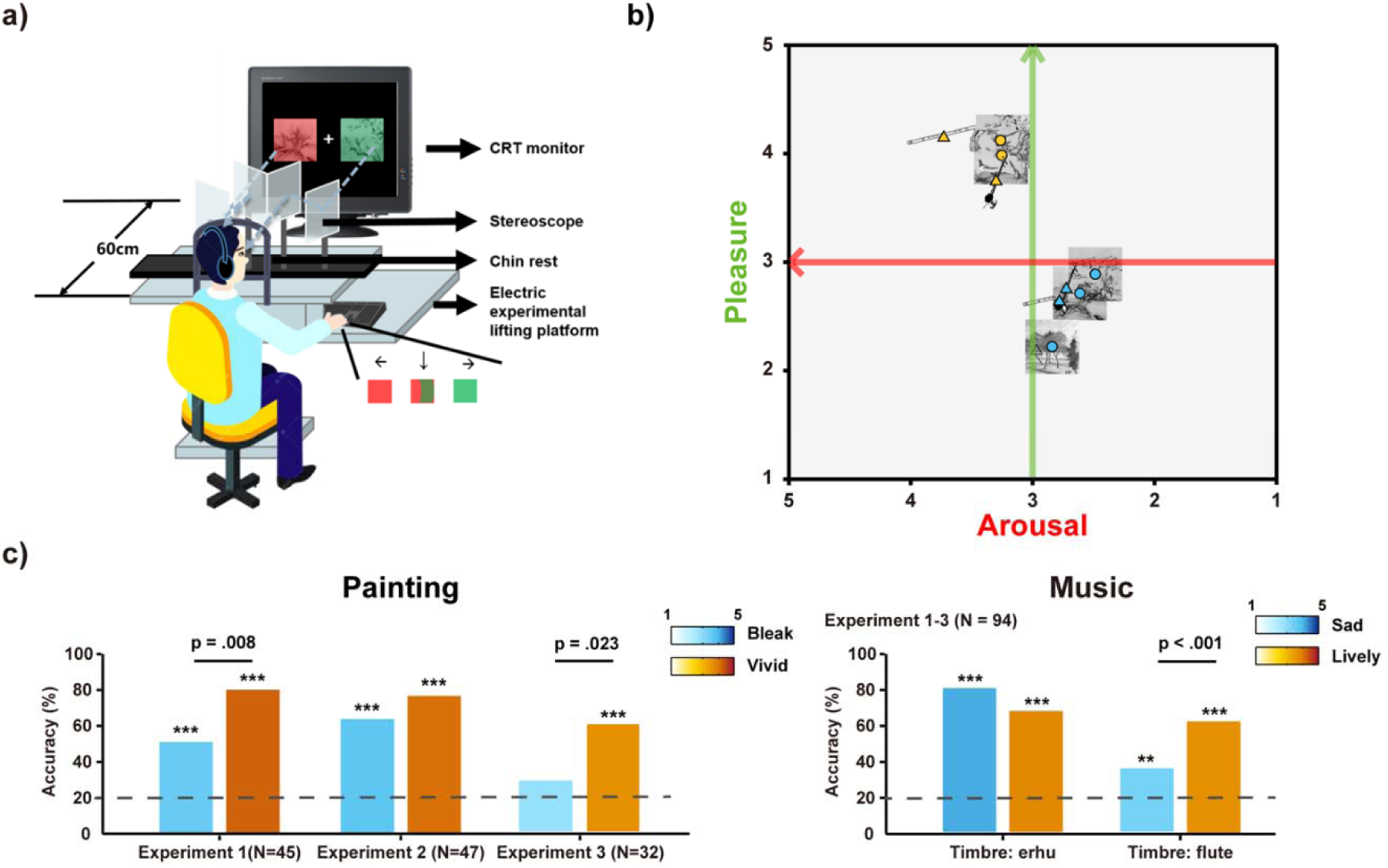
Experimental setting and emotion evaluation of stimuli. **a)** The rivalry stimuli were presented on a Cathode Ray Tube (CRT) monitor at a distance of 60 cm, while participants viewed the stimuli through the stereoscope. Participants’ head was comfortably maintained on a chin rest mounted to the electric experiment lifting platform to keep a stable visual angle. They were asked to press ‘left arrow’, ‘right arrow’ or ‘down arrow’ to indicate the color of painting (red, green or fusion) they perceived in real time. The order of buttons was counterbalanced across participants and the color of paintings was counterbalanced across trials. **b)** Subjective ratings of paintings and music show a distribution of two categories along dimensions of arousal (horizontal axis, red) and pleasure (vertical axis, green). Orange color indicates vivid paintings (circles) or lively music (triangles), blue color indicates bleak paintings (circles) or sad music (triangles). The ratings for the same painting or piece of music were pooled across Experiments 1-3. **c)** Overall identification accuracy and agreement (1 to 5 from “totally disagree” to “totally agree”, indicated by the color of bars) of the aesthetic emotions for paintings and music among all participants in Experiments 1-3. Dashed lines indicate the chance level (20%). * p <L0.05, ** p <L0.01, *** p < 0.001 by one-sample t-tests, indicating that the accuracy was significantly above chance level.

## Results

### Sad music enhanced the perceptual dominance of paintings with congruent aesthetic emotion

We calculated the average dominance duration (ADD) and predominance scores (PS) of each painting, as well as the switch rate (SR) during binocular rivalry. ADD indicates the average dominance duration, while PS refers to the proportion of accumulated dominance duration without fusion perception. SR represents the alternation rate between the two rival stimuli. We first calculated the ADD and PS for each painting type on raw data, under both background music and no music conditions. In both Experiments 1 and 2, vivid paintings exhibited significantly higher dominance than bleak paintings, as indexed by both ADD and PS scores. This effect was observed regardless of the presence of music or its aesthetic emotion and timbre (all *t*s > 2.23, FDR-corrected *p* < .031, Cohen’s *d* > 0.34, see **Figure S2** in **SI**). This consistent perceptual dominance of vivid paintings, even in the absence of music, indicates a general perceptual bias towards positive emotions. To address this, we normalized the data in subsequent analyses (see **Formula** in **Materials and Methods**) and equalized the baseline (no music condition) ADD and PS for both painting types by adjusting the luminance in Experiment 3.

As indicated by the normalized PS and ADD, sad music significantly enhanced the perceptual dominance of bleak paintings. In Experiment 1, sad music significantly increased the normalized ADD of the bleak painting compared to lively music, regardless of music timbre (erhu: *t*(39) = 2.62, FDR-corrected *p* = .006, Cohen’s *d* = 0.41, 95% CI for the mean difference = [0.04, ∞]; flute: *t*(39) = 3.05, FDR-corrected *p* =. 002, Cohen’s *d* = 0.48, 95% CI for the mean difference = [0.04, ∞], Figure 2a). This effect was replicated in Experiment 2 (*t*(42) = 2.10, FDR-corrected *p* = .021, Cohen’s *d* = 0.32, 95% CI for the mean difference = [0.02, ∞], **Figure 2b**) and in the upright but not the inverted condition of Experiment 3 (*t*(25) = 2.09, FDR-corrected *p* = .024, Cohen’s *d* = 0.41, 95% CI for the mean difference = [0.02, ∞], **Figure 2c**) when the music timbre was erhu. Moreover, in Experiment 1, when participants listened to sad music (regardless of timbre), the normalized ADD of the bleak painting significantly increased compared to the vivid painting (erhu: *t*(39) = 2.49, FDR-corrected *p* = .009, Cohen’s *d* = 0.39, 95% CI for the mean difference = [0.03, ∞]; flute: *t*(39) = 1.82, FDR-corrected *p* = .038, Cohen’s *d* = 0.29, 95% CI for the mean difference = [0.01, ∞], **Figure 2a**). However, this effect was not observed when participants listened to lively music, where the normalized ADD was equivalent between the two painting types (erhu: *t*(39) = 0.02, uncorrected p = .492; flute: *t*(39) = 1.04, uncorrected *p* = .152).

**Figure 2.**
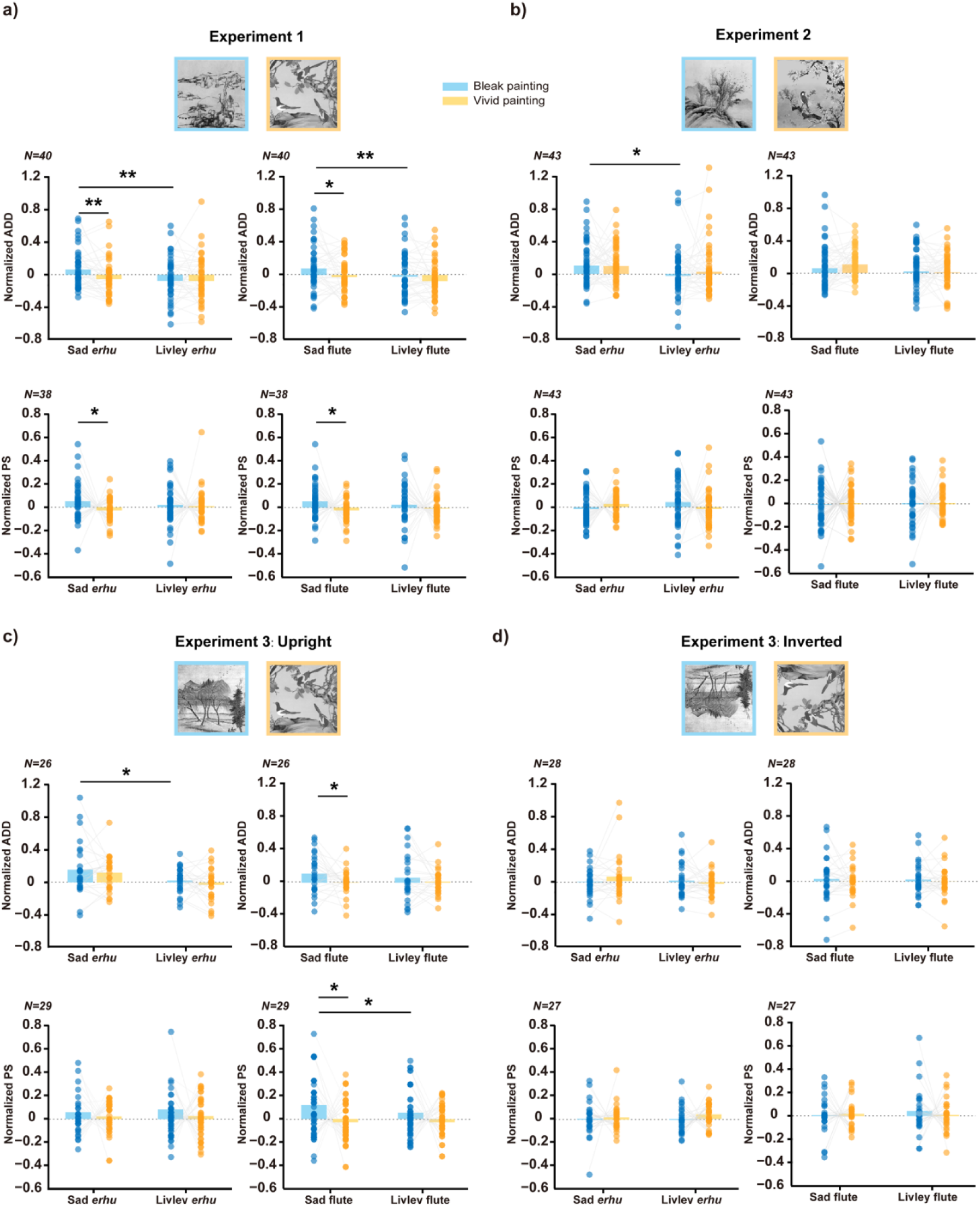
Sad music increased the normalized average dominance duration (ADD) and predominance score (PS) of upright bleak paintings. The normalized ADD and PS of upright bleak (blue) and vivid (orange) paintings were shown under sad or lively music background in Experiment 1 **(a)**, Experiment 2 **(b)** and Experiment 3 **(c)**, along with the inverted condition in Experiment 3 **(d)**. Compared to vivid paintings, sad music significantly increased the normalized ADD and PS of bleak paintings in Experiment 1 (regardless of music timbre) and Experiment 3 (especially for flute music). Compared to lively music, sad erhu music significantly increased the normalized ADD of upright bleak paintings across Experiments 1-3. These effects were not observed for inverted paintings. The valid sample sizes after excluding the outliers are shown on top of each vertical axis for each dataset. * FDR-corrected p < 0.05, ** FDR-corrected p < 0.01 by one-tailed paired t-tests.

This effect was also replicated in the upright but not the inverted condition of Experiment 3 when the music timbre was flute (*t*(25) = 1.85, FDR-corrected *p* = .038, Cohen’s *d* = 0.36, 95% CI for the mean difference = [0.01, ∞], **Figure 2c**, **2d**). These results indicate a significant top-down cross-modal congruent effect of sad music on bleak paintings.

Similarly, this congruent effect was observed in the normalized PS scores. In Experiment 1, listening to sad music significantly increased the normalized PS of the bleak painting compared to the vivid painting (*erhu: t*(37) = 1.83, FDR-corrected *p* = .037, Cohen’s *d* = 0.30, 95% CI for the mean difference = [0.01, ∞]; flute: *t*(37) = 1.83, FDR-corrected *p* = .038, Cohen’s *d* = 0.30, 95% CI for the mean difference = [0.01, ∞], **Figure 2a**). In Experiment 3, this effect persisted for upright but not inverted paintings in (**Figure 2c, d**). Specifically, when listening to sad flute music, the normalized PS of the bleak painting was significantly longer than that of the vivid painting (*t*(28) = 2.20, FDR-corrected *p* = .018, Cohen’s *d* = 0.41, 95% CI for the mean difference = [0.02, ∞]), and the normalized PS of the bleak painting was significantly longer when listening to sad flute music compared to lively flute music (*t*(28) = 2.05, FDR-corrected *p* = .025, Cohen’s *d* = 0.38, 95% CI for the mean difference = [0.01, ∞]).

### Musical tempo influenced the switch rate of both complex paintings and simple patches

We also found a significant effect of music tempo on the normalized SR in binocular rivalry. In Experiment 1, the normalized SR was significantly higher than zero when lively flute music was played (*t*(40) = 2.82, FDR-corrected *p* = .007, Cohen’s *d* = 0.44, 95% CI for the mean difference = [0.05, 0.27], **Figure 3a**), indicating that lively flute music increased the SR compared to the no-music baseline. However, sad flute music did not affect the normalized SR relative to the no-music baseline (*t*(40) = 0.83, *p* = .412, 95% CI for the mean difference = [-0.05, 0.12]). The normalized SR was also significantly faster with lively flute music compared to sad flute music (*t*(40) = 2.11, *p* = .041, Cohen’s *d* = 0.33, 95% CI for the mean difference = [0.01, 0.24]), suggesting that faster-tempo music accelerated the SR. This pattern was consistent in Experiment 2 (**Figure 3b**) and for upright but not inverted painting in Experiment 3 (**Figure 3c, d**), regardless of timbre. In Experiment 2, the normalized SR was significantly faster in the lively music condition than the no-music condition (*erhu: t*(46) = 2.84, FDR-corrected *p* = .007, Cohen’s *d* = 0.41, 95% CI for the mean difference = [0.05, 0.30]; *flute*: *t*(46) = 2.05, FDR-corrected *p* = .046, Cohen’s *d* = 0.30, 95% CI for the mean difference = [0.003, 0.29]), and in the lively music condition than the sad music condition (*erhu: t*(46) = 3.11, FDR-corrected *p* = .003, Cohen’s *d* = 0.45, 95% CI for the mean difference = [0.05, 0.30]; *flute*: *t*(46) = 2.62, FDR-corrected *p* = .012, Cohen’s *d* = 0.38, 95% CI for the mean difference = [0.04, 0.28]). In Experiment 3, the normalized SR of upright paintings was significantly faster with lively flute music compared to no music (*t*(25) = 3.61, FDR-corrected *p* = .001, Cohen’s *d* = 0.71, 95% CI for the mean difference = [0.07,0.25]), and also in the lively music condition compared to the sad music condition (*erhu: t*(25) = 2.31, FDR-corrected *p* = .030, Cohen’s *d* = 0.45, 95% CI for the mean difference = [0.01,0.18]; flute: *t*(25) = 3.11, FDR-corrected *p* = .005, Cohen’s *d* = 0.61, 95% CI for the mean difference = [0.04,0.20]).

**Figure 3.**
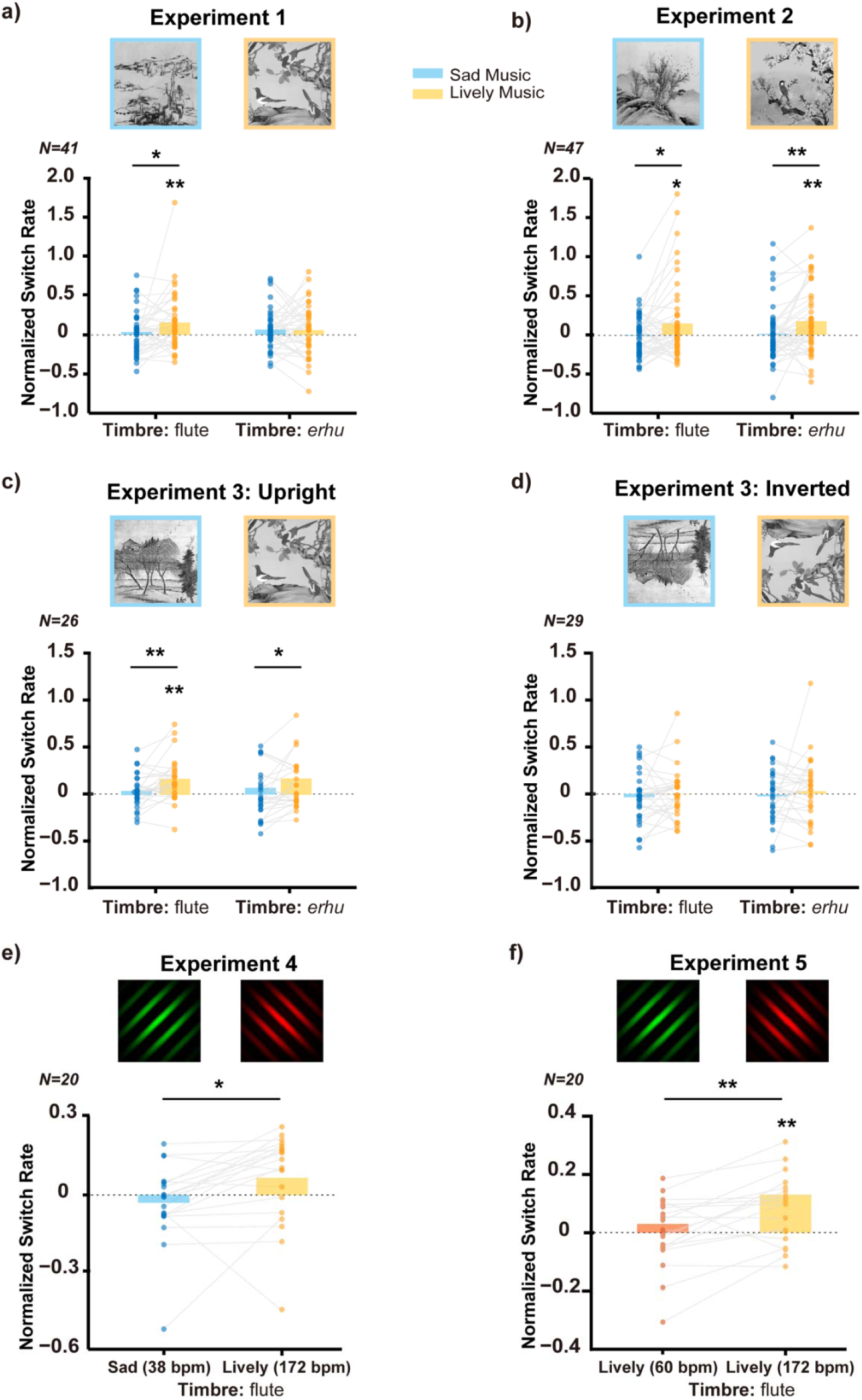
Musical tempo affected the switch rate (SR) of both paintings and patches in binocular rivalry. The normalized SR of original paintings in Experiments 1-3 (a, b, c), inverted paintings in Experiment 3 (d) and red-green patches in Experiments 4 and 5 (e, f). Compared to the sad music, listening to the lively music significantly increased the normalized SR of original paintings in both Experiments 1-3, as well as the normalized SR of patches in Experiment 4. Similarly, in Experiment 5, compared to an adjusted version of the lively music with a slower tempo (bpm = 60), the original lively music with a faster tempo (bpm = 172) significantly speeded the normalized SR of patches. The valid sample sizes after excluding the outliers are shown on top of each vertical axis for each dataset, * FDR-corrected p < 0.05, ** FDR-corrected p < 0.01 by one-sample t-tests and one-tailed paired t-tests.

To further elucidate the bottom-up property of this effect, we simplified the complex paintings into basic patches and manipulated the musical tempo in Experiments 4 and 5. In Experiment 4, to maximize the modulation of music with distinct rhythms, we chose only flute music due to its consistent modulation on the normalized SR across Experiments 1-3. With 20 new participants, we found that compared to sad flute music (38 beats per minute, bpm), lively flute music with a faster tempo (172 bpm) significantly accelerated the normalized SR of red-green patches (*t*(19) = 2.65, *p* = .016, Cohen’s *d* = 0.59, 95% CI for the mean difference = [0.02, 0.17], **Figure 3e**). In Experiment 5 with additional 20 participants, the original version of the lively flute music (172 bpm) significantly increased the normalized SR of patches compared to the no-music condition (*t*(19) = 2.38, FDR-corrected *p* = .028, Cohen’s *d* = 0.53, 95% CI for the mean difference = [0.02, 0.25], **Figure 3f**), whereas the tempo-adjusted slow version (60 bpm) did not. Furthermore, the normalized SR was significantly faster when participant listened to the original lively music compared to its slower version (*t*(19) = 2.29, *p* = .034, Cohen’s *d* = 0.59, 95% CI for the mean difference = [0.03, 0.17]). These findings indicate that musical tempo, rather than other musical features, influenced the speed of the two rivals alternated, regardless of visual complexity.

### Different correlation patterns between top-down modulation and bottom-up entrainment in sad and lively music

Having found that sad music significantly impacted top-down modulation while lively music more strongly influenced bottom-up entrainment, we further investigated the relationship of these two mechanisms, whether they are interdependent or independent. To address this, we calculated Pearson’s correlations between the normalized dominant differences (i.e. PS, ADD) of rival paintings (i.e. congruent - incongruent) and the normalized SR when listening to each music excerpt in Experiments 1-3. As shown in **Figure S3** in **SI**, when listening to sad music, the normalized SR was positively correlated with the difference in normalized PS between bleak and vivid paintings in both Experiments 1 and 2 (Exp.1: sad *erhu* music, *r* = 0.421, FDR-corrected *p* = 0.029; Exp.2: sad flute music, *r* = 0.403, FDR-corrected *p* = 0.030). However, under lively music, a significant correlation was only found in Experiment 1, and it showed a reversed pattern. Specifically, the normalized SR was negatively correlated with the difference in normalized PS between vivid and bleak paintings when listening to lively *erhu* music (*r* = -0.539, FDR-corrected *p* = 0.005), and with the difference in normalized ADD between vivid and bleak paintings when listening to lively flute music (*r* = -0.385, FDR-corrected *p* = 0.039). A similar correlation between the normalized SR and normalized PS between vivid and bleak paintings under lively flute music was found but did not survive FDR correction (*r* = -0.371, uncorrected *p* = .034). No other significant correlations were found.

### Music reward sensitivity was correlated with the modulation effect of music on binocular rivalry

Finally, we examined the relationship between these musical effects on binocular rivalry and individuals’ sensitivity to musical reward. As shown in **Table 1**, significant correlations were found with the social reward score from the Barcelona Music Reward Questionnaire (BMRQ) subsets. Specifically, the social reward score was positively correlated with the difference in normalized PS between bleak and vivid paintings when listening to sad flute music (*r* = 0.398, FDR-corrected *p* = .033), the difference in normalized ADD between vivid and bleak paintings when listening to lively *erhu* music (*r* = 0.407, FDR-corrected *p* = 0.027), and the difference in normalized ADD of the bleak painting between sad and lively *erhu* music (*r* = 0.537, FDR-corrected *p* < .001) in Experiment 2. Moreover, a higher social reward score was correlated with faster normalized SR under lively flute music (Exp. 2: *r* = 0.355, FDR-corrected *p* = .028; Exp. 1: *r* = 0.322, uncorrected *p* = .040) and sad flute music (Exp. 2: *r* = 0.394, FDR-corrected *p* = .025). These results suggest that individuals with higher sensitivity to music social reward exhibited stronger cross-modal effect of music on binocular rivalry.

**Table 1.**
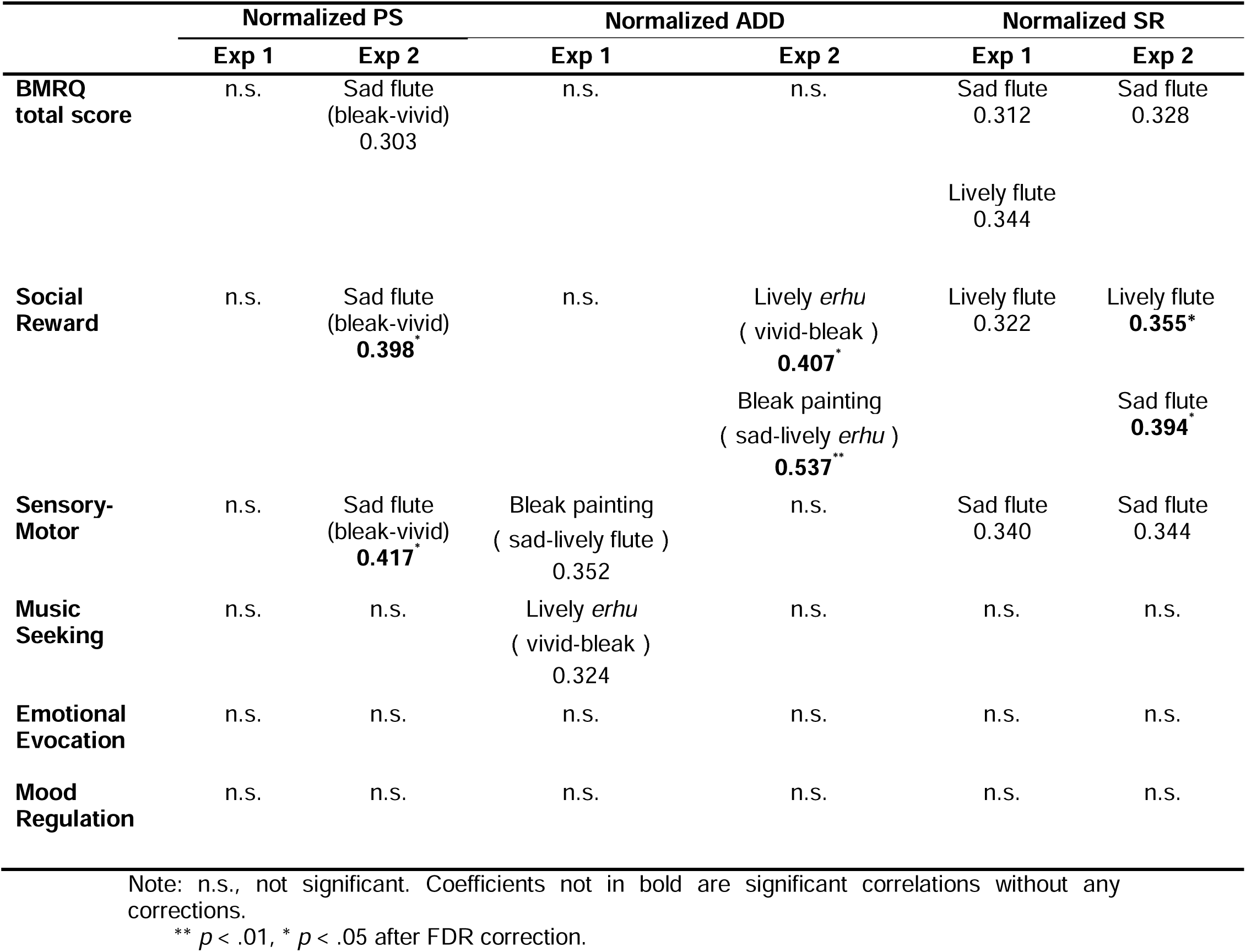
The total score and subset scores of music reward sensitivity correlated with the difference of the normalized PS and ADD between congruent and incongruent conditions and the normalized SR in binocular rivalry.

In addition, the sensory-motor subset score of BMRQ was positively correlated with the normalized PS difference between bleak and vivid paintings under sad flute music in Experiment 2 (*r* = 0.417, FDR-corrected *p* = .022). There were also significant correlations before FDR corrections between the BMRQ total score and the PS and SR effects, between the sensory-motor subset score and the ADD and SR effects, and between the music seeking subset score and the ADD effect. No other significant correlations were found.

## Discussion

In this study, we investigated how music influenced the visual perception of paintings during binocular rivalry. Our findings align with and extend previous research showing that perceptual awareness in binocular rivalry can be top-down modulated by cross-modal, higher-level mental representations (17–19). Specifically, we found that sad music facilitated the perceptual dominance of bleak paintings with congruent aesthetic emotion through top-down modulation, a process evident for upright but not inverted paintings. At a lower level of the musical hierarchy, we observed that musical tempo modulated the switch rate of visual awareness for both sophisticated paintings and simple patches in a bottom-up manner. Moreover, these top-down and bottom-up effects were interdependent and correlated with individual sensitivity to musical reward. Our findings provide evidence for predictive coding models of binocular rivalry, suggesting that music casts a hierarchical influence on the selection and alternation of visual perception in the context of visual art.

Consistent with previous studies in both experimental and real-world settings(13–15,17–23), we confirmed a cross-modal congruency effect in binocular rivalry within a novel context of artistic aesthetics. The impact of music on the perceptual awareness of paintings appears to stem from the alignment of their respective aesthetic emotions, shaped by individual prior experiences and fitting within the predictive coding framework. In this context, binocular rivalry is considered as a Bayesian perceptual inference process, where the brain chooses the hypothesis with the highest posterior probability through top-down predictions and minimizes the prediction errors from bottom-up inputs. In our study, the aesthetic emotion conveyed by music acted as a prior in shaping context and enhancing the likelihood of emotionally congruent paintings through top-down processing. This explains the prolonged dominance of congruent paintings, as they entailed less prediction errors, thus taking longer for the posterior to be destabilized and update the hypothesis. This advantage has been demonstrated in various visual forms, such as upright faces and words, even when the images are not consciously perceived (25). Notably, when the Chinese paintings were inverted, the top-down congruent effect disappeared, suggesting that this modulation relies on holistic processing, affected by the aesthetic meaning of the paintings rather than individual elements or low-level noise.

Another possible explanation is that sad music evokes a sorrowful feeling and imagery of desolate landscapes, resonating with the themes in bleak paintings. Music can evoke visual imagery and generate similar mental images among listeners from the same culture, suggesting its semantic affordances in inducing conscious congruent mental images (26,27). However, the observed musical effect cannot be fully explained at the semantic level, given the hierarchical nature of mental representations of music (9). Aesthetic emotion encompasses not only meaning but also emotional arousal and mood evocation. Therefore, the cross-modal congruency effect in binocular rivalry likely reflects a complicated top-down modulation from aesthetic emotion-related semantic representations, visual imagery and emotional arousal within a unified framework of cascaded predictive processing.

Note that the cross-modal top-down modulation was observed exclusively for sad music. The aesthetic emotion of sadness plays a dominant role in the encoding of visual mental imagery (28), which contradicts the valence-general facilitation effect found in positive and negative musical emotions (20). This discrepancy may arise from the distinction between aesthetic emotions and basic emotions in the context of art (8,26). While negative emotions are perceived as life-threatening in basic emotions (29), they are considered rewarding in aesthetic emotions (30,31). The aesthetic experience of sad music is associated with trait empathy(32), suggesting the specific role of sad music in fostering shared aesthetic emotion across modalities. Moreover, vivid paintings were more dominant than bleak ones in all conditions in Experiments 1 and 2 (see **Figure S2**), suggesting a ceiling effect where a perceptual bias driven by positive valence and high arousal persisted irrespectively of the presence of music. This bias can be partially explained by traditional interocular competition theory, which suggests that rivalry dynamics are affected by the valence and arousal of image through early bottom-up processing. The difference in visual complexity between the two types of paintings might also contribute to this effect. Notably, even after controlling for the baseline perceptual bias toward vivid paintings in Experiment 3, we still only observed the top-down modulation effect by sad music on bleak paintings. Considering that the bleak painting and sad flute music were more difficult to identify than the vivid painting and lively flute music (**Figure 1c**), respectively, the observed congruency effect for sad music but not for lively music supports the inverse effectiveness principle in multimodal integration (33).

Moreover, we uncovered a bottom-up modulation of musical tempo on the temporal alternations of binocular rivalry, a phenomenon not previously documented. While it could be contended that the effects of musical rhythm and aesthetic emotion on visual perception may coexist due to the hierarchical organization of music, our results revealed that, for patches, the accelerating effect of lively music on the normalized SR persisted when compared to sad music (Experiment 4) or a slower tempo version of lively music (Experiment 5). These findings solidify the independent role of musical tempo in influencing rivalry dynamics through bottom-up entrainment. Given the propensity of our intrinsic neural oscillations to synchronize with external rhythmic stimuli, thereby inducing subsequent perceptual oscillations (34,35), it is plausible that the oscillations of visual awareness are coupled with musical tempo, perceived as an “external” influence. Specifically, compared with sad music (0.63 Hz), lively flute music with a significantly faster tempo (2.87 Hz) has a significantly higher frequency than the frequency of visual switching in Experiments 1 to 5 (baseline SR: 0.21 Hz, 0.20 Hz, 0.25 Hz, 0.32 Hz, and 0.43 Hz, respectively, *p*s < .001, Cohen’s *d*s > 1.22, one sample t-tests), indicating audio-visual oscillation alignment. However, no visual entrainment was observed with the manipulated “lively” music with a slower tempo (1 Hz), possibly due to the restriction of intrinsic oscillations in primary auditory and visual cortices to specific δ-θ bands (2–7 Hz) in humans (36). Surprisingly, this bottom-up effect also disappeared for inverted paintings. One possible explanation is that while top-down modulation needs to act on global and meaning visual information, bottom-up entrainment may rely more on local and non-meaning information. Inverted paintings may fall between the two, resulting in a suboptimal bottom-up effect.

Additionally, we delved into the interplay between the top-down and bottom-up mechanisms unveiled in this study. Contrary to Intaitė et al. (37), who suggested that top-down and bottom-up processes independently influenced the perception of ambiguous figures, our findings revealed significant, albeit distinct, interaction patterns between these two mechanisms when participants listened to music with different aesthetic emotions. This divergence in interaction pattern may be attributed to the increased difficulty in identifying the aesthetic emotion of bleak paintings compared to vivid ones. Consequently, tracking the musical tempo became relatively easier for sad music, enabling individuals to effectively capture the hierarchical information of the music, thus facilitating the top-down congruency effect. Conversely, enhanced entrainment to lively music resulted in a reduced cross-modal congruency effect in Experiment 1. This can be explained by the inherent perceptual dominance advantage of vivid paintings, whereby better synchronization with lively music led to faster switching, ultimately diminishing the cross-modal congruency effect.

Interestingly, the top-down modulation and bottom-up entrainment mechanisms were not only interdependent but also associated with individual music reward sensitivity, especially in the social reward sub-dimension. This finding is consistent with previous studies showing that social interaction ability was correlated with the perceptual switch of red and green checkerboards (38) but did not affect the awareness of rival gratings in a clock-wise rotating predictive context (39). It is plausible that as individuals experience greater social bonding through music, the capacity of music to forge connections with the visual world and foster shared aesthetic perception becomes more pronounced. The lack of correlations between top-down and bottom-up mechanisms and their relationships with music reward sensitivity in Experiment 3 may be caused by the relatively small sample size.

The limitations of the current study include imperfect control over the visual complexity of paintings and auditory complexity of music. Due to the limited numbers of paintings and pieces of music used, and slightly inconsistent result patterns across stimuli and experiments, the generalization of our findings needs further investigations with larger sample sizes. Additionally, although aesthetic emotion is a universal term in art, we cannot isolate the independent roles of musical emotion and semantics in binocular rivalry of paintings. It is also unclear at which level the interaction of aesthetic emotions in music and painting occurs. Future studies should determine the specific level and corresponding neural mechanisms underlying cross-modal interactions in audio-visual art scenes.

In conclusion, our study revealed two noteworthy cross-modal mechanisms on binocular rivalry in an artistic context. The hierarchical organization of music, specifically its aesthetic emotion and tempo, modulated the selection and alternation of visual awareness of paintings, respectively. Notably, these two mechanisms were interdependent and correlated with individual sensitivity to music social reward. These findings support the predictive coding models in explaining binocular rivalry (40,41) and deepen our understanding of the complex cognitive processes underlying multisensory perception. By revealing how music can enhance the appreciation and perception of abstract visual works in an immersive and multisensory manner, our findings pave the way for future research aimed at exploring the potential applications of music in art education and therapy.

## Materials and Methods

### Participants

A total of 138 naïve observers participated in the 5 main experiments. Forty-five subjects participated in Experiment 1 (26 females, mean age = 22.7 ± 3.0 years), 49 participated in Experiment 2 (28 females, mean age = 23.2 ± 2.7 years), 32 participated in Experiment 3 (19 females, mean age = 24.4 ± 2.8 years), 20 participated in Experiment 4 (10 females, mean age = 22.5 ± 3.1 years) and the other 20 participated in Experiment 5 (11 females, mean age = 25.3 ± 3.5 years), with 28 (14 females, mean age = 22.5 ± 2.5 years) participated in both Experiments 1 and 2. A power analysis using G*Power (42) revealed that a sample size of at least 25 participants and 14 participants could ensure sufficient power (one-tailed; Cohen’s d = 0.7; α = 0.05; 1-β = 0.99) to detect a congruency effect in cross-modal binocular rivalry using images (17) and patches (16), respectively. Given that the traditional Chinese paintings used in Experiments 1-3 were complex images which were never addressed in previous studies, we increased the sample size to over thirty. All participants were screened using the Beck Depression Inventory— Second Edition (BDI-II) (43) to exclude those with potential impact of depression (BDI score < 4), and completed the Chinese version of Barcelona Music Reward Questionnaire (BMRQ) (44) to assess music reward sensitivity. All participants had normal hearing (binaural pure tone average < 25 dB hearing level at 250-8000 Hz), normal or corrected-to-normal vision, normal stereoacuity, and no strong eye dominance as defined by the binocular rivalry calibration before the formal experiments. Neither color blindness nor color weakness were reported from all participants. The research was approved by the Ethics Committee of the Institute of Psychology, Chinese Academy of Sciences. All participants provided written informed consent before the study and were paid 50 RMB per hour for their participation.

### Apparatus

The rival stimuli were presented using Psychtoolbox-3 (http://psychtoolbox.org/) in MATLAB R2016a (MathWorks) on a 17-inch CRT monitor (Lenovo LXB-HF769A, 1024×768 resolution, 85-Hz refresh rate) at a distance of 60 cm in all experiments. After the participants pressed the space button of the keyboard, two rival stimuli were displayed side by side on the CRT monitor and fused with a mirror stereoscope. A chinrest mounted to the electric lifting platform was used to maintain the head position. The auditory stimuli were initialized by PsychPortAudio and presented binaurally through a Sennheiser HD650 Pro headphone.

### Visual Stimuli

In Experiments 1-3, the visual stimuli were paintings selected from a database for emotion and aesthetic analysis of traditional Chinese paintings which was validated by 20 participants without professional art training (45). These Chinese paintings were cropped into squares with good picture clarity, suitable size, and arrangement, sharing the same size and resolution (100*100 pixel for pretest and 354*354 pixel for binocular rivalry tasks). To minimize the “semantic loss” caused by cropping, only the seals, inscription or other blank areas were cropped. The color of each painting was rescaled into gray and other low-level properties such as luminance and contrast were controlled. An exception is that, in Experiment 3, the luminance of the bleak painting was increased to eliminate the perceptual advantage of the vivid painting, ensuring an equal predominance score under the no-music baseline condition. All image editing operations were conducted using Adobe Photoshop CC 2017 (Adobe Systems Incorporated). The final five Chinese traditional paintings (2 vivid and 3 bleak, one vivid painting was replicated in Experiment 3, see **Figure 2**) were selected by matching the aesthetic emotion (vivid or bleak) with chosen music excerpts (see **Figure S1** and **Pretest** for details). The aesthetic emotion of paintings was confirmed by participants after the binocular rivalry task (see **Figure 1c** and **Subjective assessments** for details). As shown in **Figure 1b**, the vivid and bleak paintings differed significantly on the level of pleasure (*t*(93) = 11.92, *p* < .001, Cohen’s *d* = 1.23, 95% confidence interval (CI) for the mean difference = [1.04,1.45]), and arousal (*t*(93) = 4.82, *p* < .001, Cohen’s *d* = 0.50, 95% CI for the mean difference = [0.36,0.86]). To minimize binocular fusion perception, the tone of the paintings was adjusted to red or green, and the tone was counterbalanced across trials for each pair of paintings. In Experiments 4 and 5, the visual stimuli were one red and one green Gabor patches of sine-wave gratings (5 cycles/deg) with orthogonal orientations (±45° from vertical), smoothed at the edges with a Gaussian filter.

### Auditory Stimuli

The auditory stimuli were four 1-minute Chinese folk music clips with combinations of aesthetic emotions (sad or lively) and timbres (flute or *erhu*), chosen from 8 music excerpts through an audio-visual matching pretest (**Figure S1**, see **Pretest** for details).

The two chosen sad music clips are “Performing with an only string” 《独弦操》 (sad *erhu* music), “Solitary Orchid Greeting the Spring” 《幽兰逢春》 (sad flue music), and the two lively music clips are “Birds Are Singing in Tranquil Valley”《空山鸟语》 (lively *erhu* music), “Spring Visiting Xiang Jiang River”《春到湘江》 (lively flute music). The aesthetic emotions of music were confirmed by participants in Experiments 1-3 after the binocular rivalry task (see **Figure 1c** and **Subjective assessments** for details). As shown in **Figure 1b**, the lively and sad music differed significantly on pleasure (*t*(123) = 28.29, *p* < .001, Cohen’s *d* = 2.54, 95% CI for mean difference = [0.78, 1.21]) and arousal (*t*(123) = 3.70, *p* < .001, Cohen’s *d* = 0.33, 95% CI for mean difference = [0.19, 0.64]). The average tempo of music excerpts was 39 beats per minute (bpm) for sad *erhu* music, 38 bpm for sad flute music, 126 bpm for lively *erhu* music, and 172 bpm for lively flute music. In Experiment 5, the lively flute music (172 bpm) was modified into a slower version by only changing the musical tempo into 60 bpm. The music excerpts were 32-bit at a sample rate of 44,100 Hz and matched in terms of average root mean square sound pressure level (SPL). The intensity levels of auditory stimuli were calibrated in the experimental environment using a Larson-Davis sound level meter (Model 831, Depew, NY), and fixed at 65 dB SPL in all experiments.

### Procedure

In Experiments 1-3, the five auditory conditions (no music baseline, lively flute, sad flute, lively erhu, sad erhu) were randomly presented in a total of 40 trials, with 8 trials for each condition (In Experiment 3, upright and inverted paintings were presented for 4 trials each). In each experiment, only one pair of the rival (vivid and bleak) paintings was presented, with counterbalanced tone (red and green) and left-right visual field across trials within subject. Before the task, calibration was completed after participants adjusted the stereoscope to ensure that the two images were perfectly fused and no strong dominance of one eye was reported. In each one-minute trial, participants started by pressing the ‘space’ key on the computer keyboard, and then two rival paintings were presented to each eye field while background music played. They were instructed to press the left, right or down key (see Figure 1a) to report the color (red, green or mixed) of painting they perceived, and release the key immediately after pressing it to record their rivalry fluctuations. The order of keys was counterbalanced across participants.

In Experiments 4 and 5, participants completed 24 one-minute trials with three randomly played auditory conditions (no music baseline, lively flute, sad flute in Experiment 4; no music baseline, original lively flute, slower tempo lively flute in Experiment 5). The procedure was the same as that in Experiments 1-3, and participants were asked to track their perception of gratings’ color (red or green).

### Pretest

To select the music excerpts and paintings with the best matched aesthetic emotions, additional 17 subjects (10 females, mean age = 23.1 ± 2.0 years) who have not participated any main experiment took part in the pretest for stimuli selection. Specifically, a circular arena with a dot in the center position was displayed on a computer screen. Initially, 23 traditional Chinese paintings (see Figure S1a in SI) from a validated Chinese paintings database (45) were randomly presented outside and around the arena (see Figure S1b in SI). Participants were informed that the distance between the final position of painting and the center of arena indicates the congruence of aesthetic emotion between the painting and the music. While listening to music, they were instructed to move any item to the proper position by “dragging-and-dropping” it and click the “Done” button to confirm their final decision to turn to the next trial. In each trial, the same music was played in a loop until the participant completed the final arrangement. To increase the consistency of evaluation, the same song was played three times consecutively. This method was established to obtain similarity data from multiple item arrangements (46,47) and recently has been widely used to get the similarity of the object features (48), semantic distance (49) and social distance (50).

The similarity of aesthetic emotion between music and painting was calculated using Euclidean distance. We averaged the distances between the center of arena and the positions of paintings across the three trials under the same music to ensure the stability of the evaluation results. For each music, the four paintings with the smallest distance were considered as the best-matched. The final four paintings that were most frequently selected as the best-matched ones by all participants were used in the rivalry tasks. As shown in Figure S1c in SI, for sad music, 58.8%, 41.2% and 52.9% of participants chose the final three bleak paintings for “Performing with an only string”, and 41.2%, 47.1% and 41.2% of participants chose the final three bleak paintings for “Solitary Orchid Greeting the Spring”. For lively music, 88.2% and 11.8% of participants chose the final two vivid paintings for “Birds Are Singing in Tranquil Valley”, and 52.9% and 88.2% of participants chosen the final two vivid paintings for “Spring Visiting Xiang Jiang River”. Notably, although only 11.8% of participants matched one of the vivid paintings with “Birds Are Singing in Tranquil Valley”, we decided to include it in the main experiments as it shares elements (i.e. birds and trees) and spatial layout with the other vivid painting. This allowed us to maintain the consistency in the semantic content of diverse visual stimuli and minimize fusion perception.

### Subjective assessments

After the binocular rivalry task in Experiments 1-3, participants further assessed the music and paintings they already heard and seen in the rivalry task. They were asked to rate the pleasure (from 1 “negative” to 5 “positive”) and arousal (from 1 “calm” to 5 “aroused”) as well as choose one of five alternatives to represent the given aesthetic emotion. For paintings, the alternatives were “vivid”(“生机”), “elegant”(“雅致”), “magnificent”(“气势”), “quiet”(“清幽”) and “bleak”(“萧瑟”), while the options for music were “lively” (“灵动”), “excited” (“激扬”), “soft” (“轻柔”), “deep” (“深沉”) and “sad” (“悲伤”). All participants expressed their agreement on the given aesthetic emotion using a Likert scale ranging from 1 “totally disagree” to 5 “totally agree”, regardless of whether they could accurately identify the aesthetic emotion.

We focused on the identification accuracy, agreement of given aesthetic emotion, and the pleasure-arousal rating for each music and painting. The identification accuracy was considered as the proportion of people that successfully identified the given aesthetic emotion among all participants. One-sample t-tests were performed to examine if the identification accuracy was substantially different from the chance level (20%). The agreement of a specific aesthetic emotion was averaged across subjects for each painting and music piece, with the higher agreement indicating stronger participant consistency for that aesthetic emotion. For each pair of paintings or each pair of music, paired t test was utilized to compare the identification accuracy, agreement and pleasure-arousal rating between diverse aesthetic emotions.

For paintings, as illustrated in **Figure 1c (left)**, the identification accuracy of the vivid and bleak paintings was significantly higher than chance level (20%) in Experiments 1 and 2 (*p*s < .001, Cohen’s *d*s> 0.76, by one-tail one sample t-test, 0.80 ± 0.40 and 0.77 ± 0.42 for the two vivid paintings, 0.51 ± 0.50 and 0.64 ± 0.48 for the two bleak paintings). However, in Experiment 3, only the vivid painting utilized in Experiment 1 was significantly identifiable (0.59 ± 0.50, *t*(44) = 9.95, *p* < .001, Cohen’s *d* = 1.48, 95% CI for the mean difference = [0.70, ∞]), while the identification accuracy of the bleak painting did not differ significantly from the chance level (0.28 ± 0.46, *t*(31) = 1.01, *p* = .161). Compared with vivid aesthetic emotion, participants found it harder to discriminate bleak aesthetic emotion from other four alternatives for the first pair of paintings in Experiments 1 (*t*(44) = 2.79, *p* = .008, Cohen’s *d* = 0.42, by two-tailed paired t-test, 95% CI for the mean difference = [0.08, 0.50]) and 3 (*t*(31) = 2.40, *p* = .023, Cohen’s *d* = 0.42, 95% CI for the mean difference = [0.05, 0.58]), with no difference between the second pair of paintings in Experiment 2 (*t*(46) = 1.43, *p* = .160, Cohen’s *d* = 0.21, 95% CI for the mean difference = [- 0.05,0.31]). There was no significant difference in degree of agreement on the given aesthetic emotion (bleak or vivid) between the two groups of participants in Experiments 1 and 2 (*p*s > .05, Cohen’s *d*s < 0.20). However, the degree of agreement was significantly higher in vivid paintings than in bleak ones in both Experiments 1 (*t*(44) = 4.11, *p* < .001, Cohen’s *d* = 0.61, 95% CI for mean difference = [0.34,0.99]), 2 (*t*(46) = 2.70, *p* = .010, Cohen’s *d* = 0.39, 95% CI for mean difference = [0.09,0.63]) and 3 (*t*(31) = 3.54, *p* = .001, Cohen’s *d* = 0.63, 95% CI for the mean difference = [0.36, 1.33]).

For music, as shown in **Figure 1c (right)**, the identification accuracy of lively and sad music was significantly higher than chance level (20%) (*p*s < .01, 0.67 ± 0.47 and 0.62 ± 0.49 for two lively music, 0.80 ± 0.40 and 0.35 ± 0.48 for two sad music in Experiment 1-3). In Experiment 1-3, when the timbre was *erhu*, there was no significant difference between the identification accuracy of lively music and sad music, whereas sad aesthetic emotion was more difficult to be perceived than lively aesthetic emotion when the timbre was flute (*t*(93) = -3.91, *p* < .001, Cohen’s *d* = 0.40, 95% CI for mean difference = [-0.40, -0.13]). Consistently, the degree of agreement was significant lower in sad music than in lively music regardless of timbre (flute: *t*(93) = -3.13, *p* = .002, Cohen’s *d* = 0.32, 95% CI for mean difference = [-0.56, -0.12]); *erhu*: *t*(93) = -2.59, *p* = .011, Cohen’s *d* = 0.27, 95% CI for mean difference = [-0.53, -0.06]).

### Data analysis

We calculated the predominance scores (PS) and average dominance duration (ADD) of each painting, as well as the switch rate (SR) during binocular rivalry. To uncover the specific effects of musical, all measures were normalized to the no-music baseline using the formula below.

The PS, ADD and SR in binocular rivalry were normalized using the following formula:

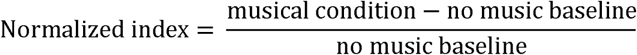

PS refers to the proportion of cumulate dominance duration (d) of each painting in the total duration without fusion perception during rivalry. In each trial, the bleak painting and vivid painting were calculated separately using the following formula:

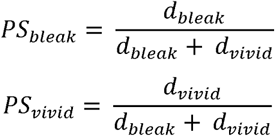

ADD indicates the average duration across the total number (N) perceived by the participant, and calculated in each trial as:

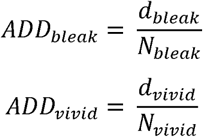

SR represents the alternative rate between the two paintings in Experiments 1 and 2 or between the two patches in Experiments 3 and 4, and this index was calculated across trials as:

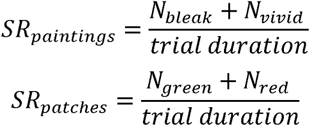

For each normalized measure in each musical condition, outliers (2.5 standard deviation away from the mean) were first excluded, and one-sample t-tests were first performed to examine the specific musical effect. Then one-tailed paired t-tests were performed on the normalized PS and ADD between congruent and incongruent music-painting pairs to test the facilitating effect of cross-modal congruency effect in Experiments 1-3. Furthermore, two-tailed paired t-tests were performed in all experiments to compare the normalized SR under each musical condition. Pearson’s correlation was used to test the relationship between the top-down (normalized PS and ADD) and bottom-up (normalized SR) effects within the same musical condition. Finally, Pearson’s correlations were calculated between the BMRQ score and normalized measures to test the relationship between musical reward sensitivity and musical effects on binocular rivalry. False discovery rate (FDR) correction was used for multiple comparisons.

## Author Contributions

Conceptualization: Y. Du and X.T. Jiang; Funding acquisition: Y. Du; Formal Analysis: X.T. Jiang and Y.H. Cheng; Methodology: X.T. Jiang, Y.H. Cheng, Y. Jiang and Y. Du; Investigation: X.T. Jiang and Y.H. Cheng; Visualization: X.T. Jiang and Y. Du; Supervision: Y. Jiang and Y. Du; Writing (original draft): X.T. Jiang; Writing (review and editing): X.T. Jiang, Y.H. Cheng, Y. Jiang, and Y. Du.

## Acknowledgments

This research was supported by grants from the STI 2030-Major Projects 2021ZD0201500 (Y.D.) and 2021ZD0203800 (Y.J.), the National Natural Science Foundation of China 32430043 (Y.J.), and the Strategic Priority Research Program of Chinese Academy of Sciences XDB32010300 (Y.D. and Y.J.). The authors would like to thank Yan Gao for providing the painting dataset, Baishen Liang and Xiaonan Li for help with pretest preparation, and Peijun Yuan and Tian Yuan for setting up binocular rivalry task.

## Supplemental Information (SI)

**Figure S1.**
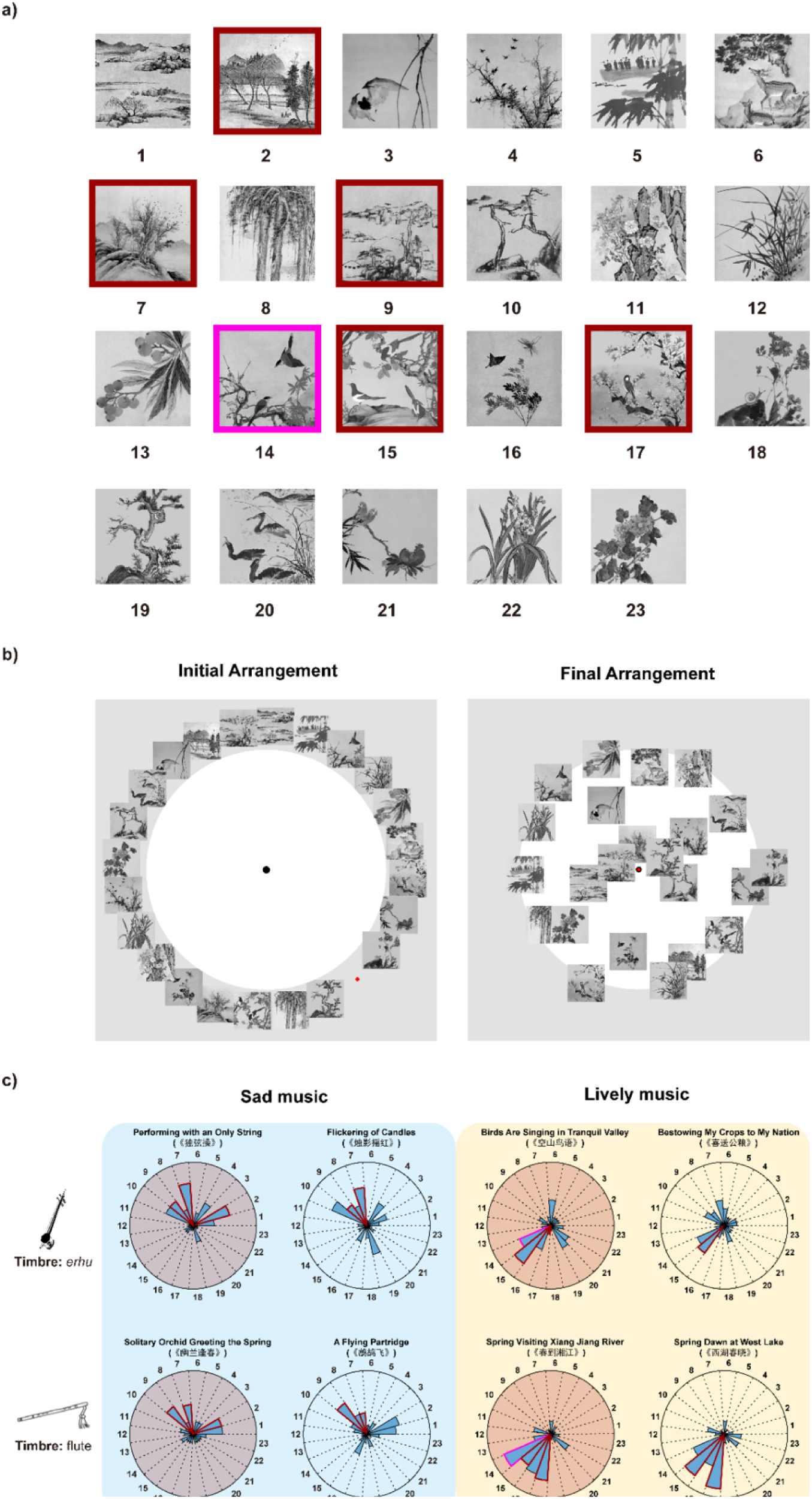
Music-painting matching pretest (N = 17). a) The original 23 traditional Chinese paintings used in the pretest. b) Procedure of the pretest. The paintings were initially presented randomly outside an arena with music playing (left panel). The participants were instructed to drag the paintings inside the arena according to the similarity of the aesthetic emotion they felt between the painting and the current music (closer to the center of the arena for more similarity). The right panel shows an example of the final arrangement when listening to sad erhu music. c) The proportion of each painting that was chosen as the closest four to each of the 8 music excerpts. In a) and c), the red borders indicate the final selected paintings used as the rivalry stimuli in the main experiments, and the pink borders indicate those with high similarity to the music excerpt but were not used because of their high rate of fusion or obvious perceptual dominance of vivid painting during binocular rivalry.

**Figure S2.**
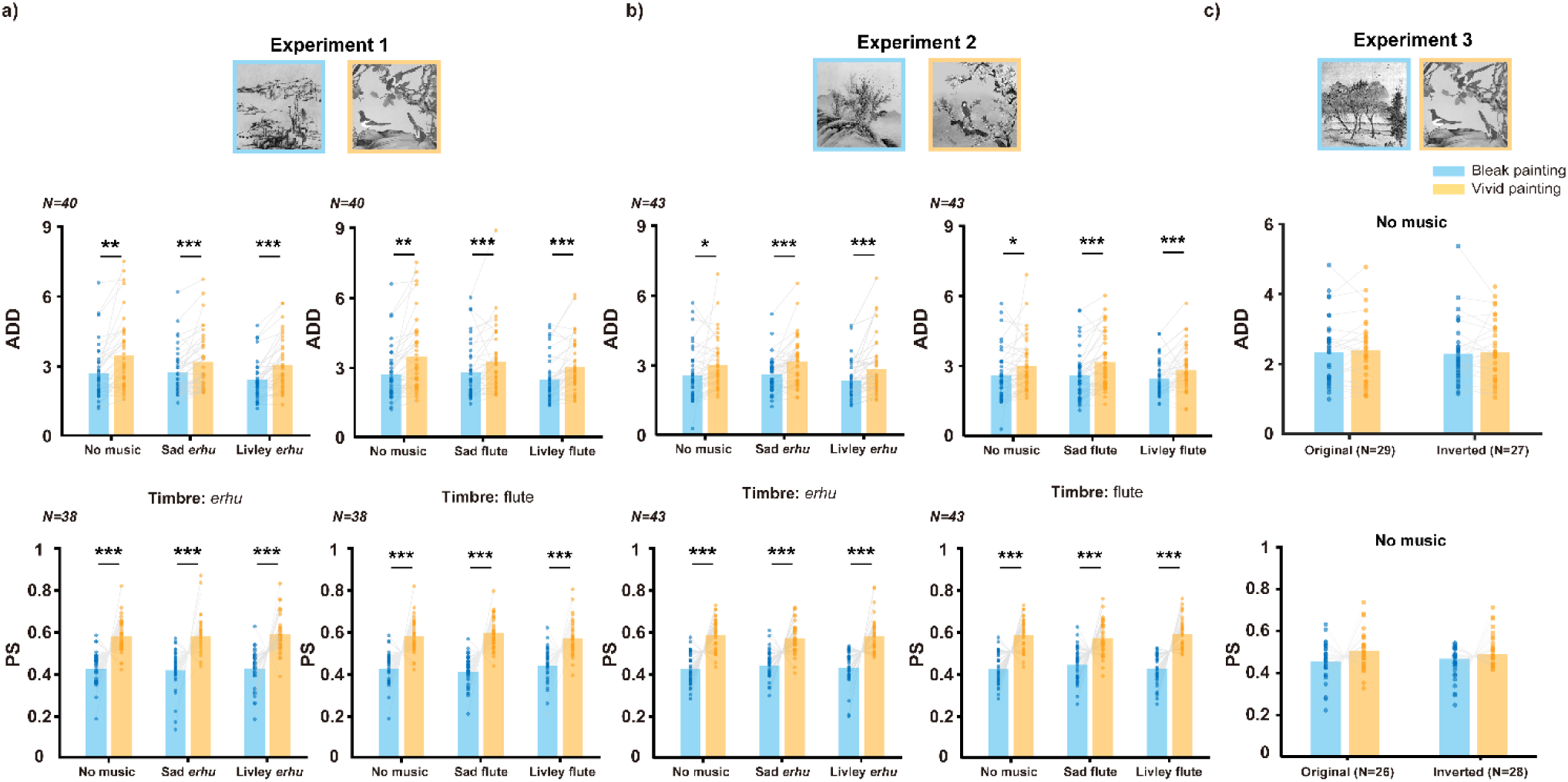
Perceptual dominance of vivid paintings regardless of music in Experiments 1 and 2, and equal dominance of bleak and vivid paintings in no music baseline in Experiment 3. In both sad and lively musical conditions and the no music baseline condition, the average dominance duration (a) and the predominance score (b) of the vivid painting were significantly higher than those of the bleak painting in both Experiments 1 and 2. (c) In no music baseline condition, there is no significant difference in the predominance score and the averaged dominance duration of bleak and vivid paintings. * p < 0.05, ** p < 0.01, *** p < 0.001 by paired t-tests, FDR corrected.

**Figure S3.**
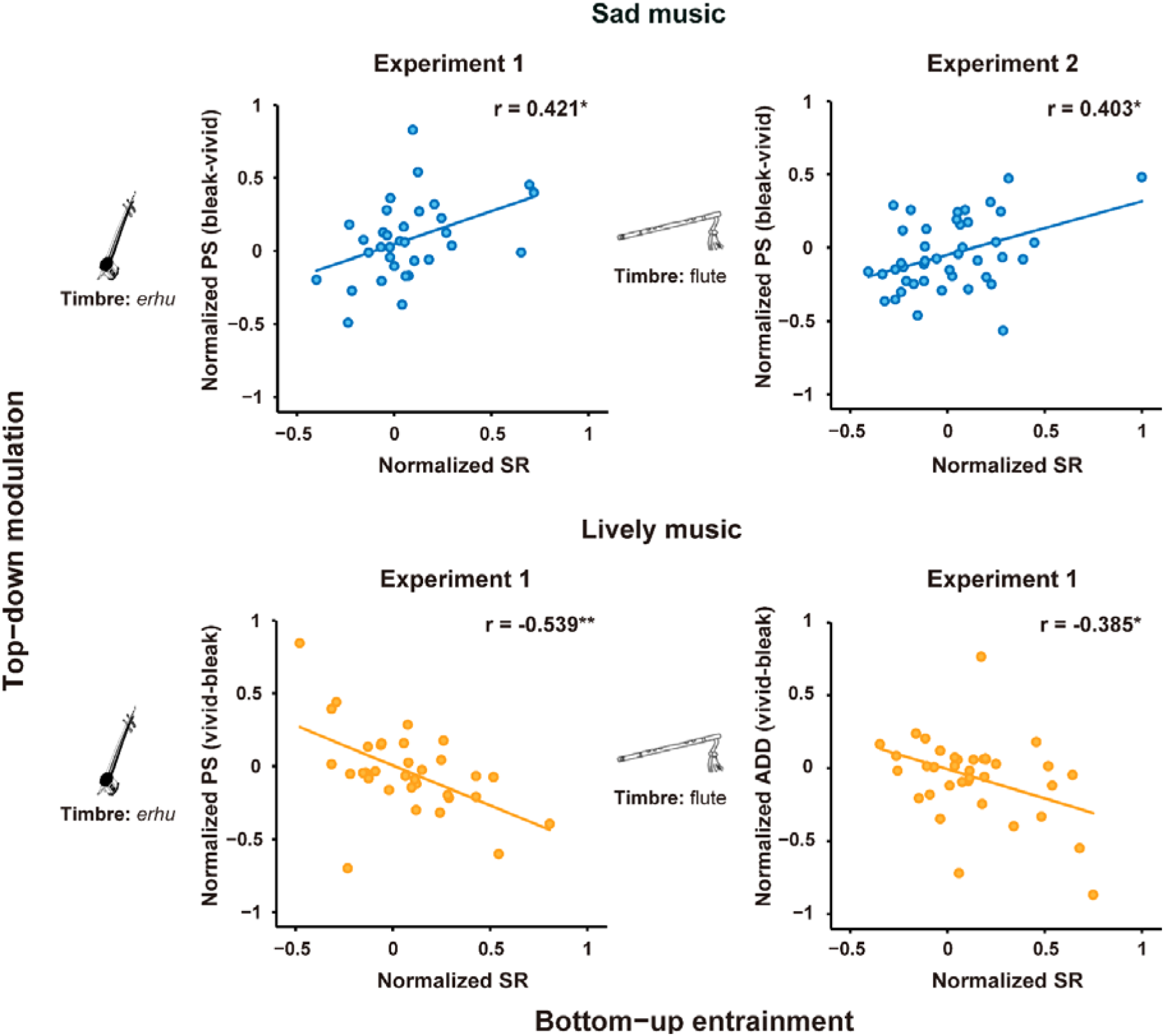
Different correlation patterns between top-down modulation (indexed by PS and ADD) and bottom-up entrainment (indexed by SR) in sad and lively music. The correlations of the dominant difference (indexed by normalized PS) between bleak and vivid paintings and the normalized SR when listening to sad music (blue points) in Experiment 1(top left panel) and 2 (top right panel). The correlations of the dominant difference (bottom left panel: indexed by normalized PS, bottom right panel: indexed by normalized ADD) between vivid and bleak paintings and the normalized SR when listening to lively music (orange points) in Experiment 1. When listening to sad music, the normalized SR was positively correlated with the difference of normalized PS of bleak and vivid painting. Reversely, when listening to lively music, the normalized SR was negatively correlated with the difference of normalized PS of bleak and vivid painting. The statistics were conducted using Pearson’s correlations, and fitted by linear model. * FDR-corrected *p* < 0.05, ** FDR-corrected *p* < 0.01.

